# Physics of Lumen growth

**DOI:** 10.1101/294264

**Authors:** Sabyasachi Dasgupta, Kapish Gupta, Yue Zhang, Virgile Viasnoff, Jacques Prost

**Affiliations:** Mechanobiology Institute, National University of Singapore, Singapore 117411, Singapore; Laboratoire Physico Chimie Curie, Institut Curie, PSL Research University, CNRS UMR168, 75005 Paris, France; CNRS UMI3639, Singapore 117411, Singapore; Department of Biological Sciences, National University of Singapore, Singapore 117411, Singapore

**Keywords:** osmoregulation, membrane pumps, lumens, tissue mechanics

## Abstract

We model the dynamics of formation of intercellular secretory lumens. Using conservation laws, we quantitatively study the balance between paracellular leaks and the build-up of osmotic pressure in the lumen. Our model predicts a critical pumping threshold to expand stable lumens. Consistently with experimental observations in bile canaliculi, the model also describes a transition between a monotonous and oscillatory regime during luminogenesis as a function of ion and water transport parameters. We finally discuss the possible importance of regulation of paracellular leaks in intercellular tubulogenesis.

---

Epithelial lumens are ubiquitous in organs. They originate from cavities or tubes surrounded by one (seamless lumen) or multiple cells (1). Ions and other bioactive molecules are secreted into the cavities and, if the lumen is open, flow with the physiological medium. The creation of the lumens orginates from several classes of morphogenetic events (1). In the case of closed lumens (such as acini, blastocytes, canaliculi), ion secretion into the forming cavity creates an osmotic pressure. This results in the passive transport of water into the lumen (most often mediated by aquaporins), which constitutes a major driving component for lumen expansion. This osmotic pressure hypothesis was experimentally proposed in the 1960s (2-4). The expansion is mechanically restrained by periluminal tension. In the case of multicellular lumens (eg: cysts (5-7)), tension results from the contraction of the cells surrounding the lumen. In the case of the intercellular domain, the tension arises from the cortical actin layer surrounding the cavity (8).

Fig. 1a illustrates a lumen separating adjacent membranes between two primary rat hepatocytes (liver cells). The contact area between both cells presents an intercellular cleft of around 30-50 *nm* (9) that accommodates transcellular proteins, adhesion proteins and peptidoglycans. The development of the lumen occurs within 5 to 6 hours. *In vivo,* closed lumens eventually merge into a network of tubules called canaliculi (2*µm* diameter and 500 *µm* long). We recently showed that the shape of these lumen is controlled by the balance of osmotic pressure and anisotropic cortical tension (10). Hepatocyte doublets can be used as meaningful simplified surrogates to study lumen formation (8, 11, 12). In this instance functional canaliculi grow as spherical caps spanning part of the intercellular space. The simple geometry of the system constitutes an appealing case for quantitative studies.

**Fig. 1.**
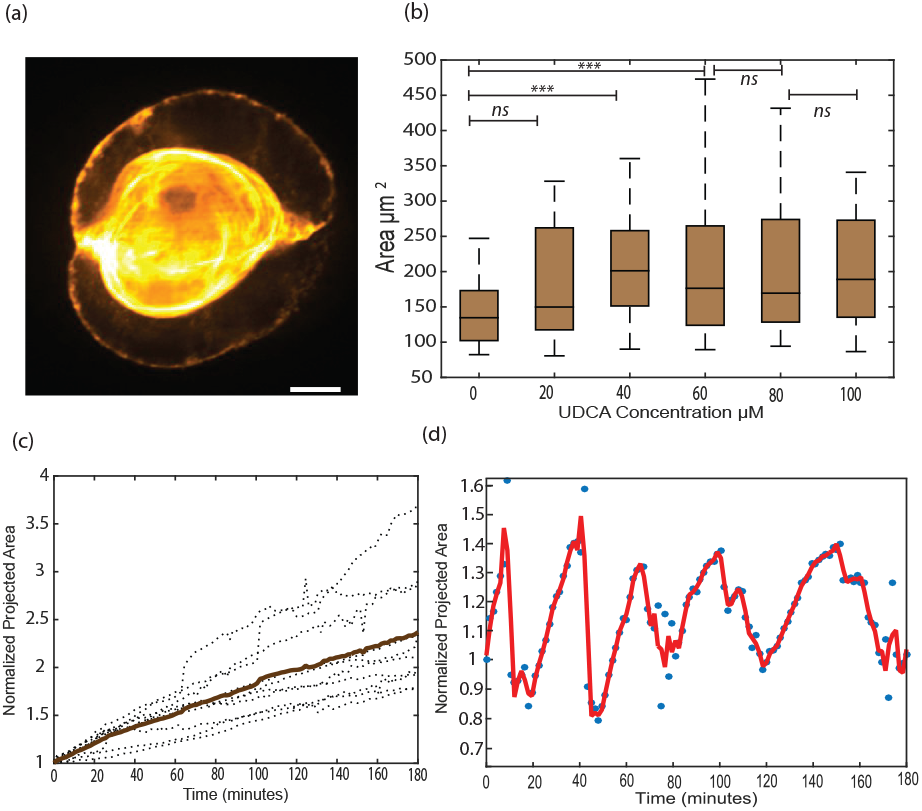
a:Structured illumination image of a typical bile canaliculi creating a lumen between two hepatocytes (scale bar 2 *µ*m). b: Increase of projected area of canaliculi at steady state upon continuous bile secretion stimulation by different dose of UDCA (Ursodeoxycholic acid) (n=20 for each dose).c: Linear growth of the canaliculi (dotted line:individual cell, bold line:average) under reduced contractility condition (1 *µ*M blebbistatin) d: Sustained Oscillatory dynamics under native contractility conditions. Bile canaliculi projected area are normalized by their size at t=0.

However, this process is rather generic for many kinds of lumen such as Ciona Notochord lumen (1, 13, 14) or kidney lumens(15). Fig. 1b-c also shows that the steady shape of the lumen depends on the secretory activity, which is boosted by the addition of Ursodeoxycholic acid (UDCA). The growth of the lumen can either be monotonous (Fig. 1c) or pulsatile (Fig. 1d) depending on the periluminal tension and secretory activity. A steady secretion in a closed lumen implies the concomitant existence of leakage. Its nature is likely paracellular (through the nanometer cleft between cells). In the case of multicellular lumen, a few models and experimental studies have considered the role of leaks (originating either from the rupture of cell-cell contacts (7) or permeation across the endothelial layer (16)) during the growth of the lumen. For intercellular lumens, however, the morphogenetic consequences of the leak modulation by the paracellular cleft property have hardly been investigated, either experimentally or theoretically.

Here, we provide a theoretical quantitative study on the balance between secretory activity, leak and mechanics that determines canaliculi nucleation and growth. Our minimalistic description of lumen expansion identifies the physiologically relevant range of parameters required to establish a stable intracellular cavity and dictate its dynamical properties.

## Modeling Assumption

We consider the lumen as two symmetrical contractile spherical caps (Fig. 2) with a radius of curvature R and a contact angle *θ* at the lumen edge. The lumen elongates parallel to the cell-cell contact over a distance *r*_*l*_ and its apex height is *h*. The remaining paracellular adhesive cleft has a thickness e. As the lumen develops, the dimension of the spherical caps vary but the cell contact remains fixed with a total size L. We established the expressions of the conservation laws in the lumen and in the cleft accounting for this geometry. All results are in the scaled units of the model (See SI appendix, Table S1) as well as in “international units” based on the estimations derived in SI Appendix (2).

**Fig. 2.**
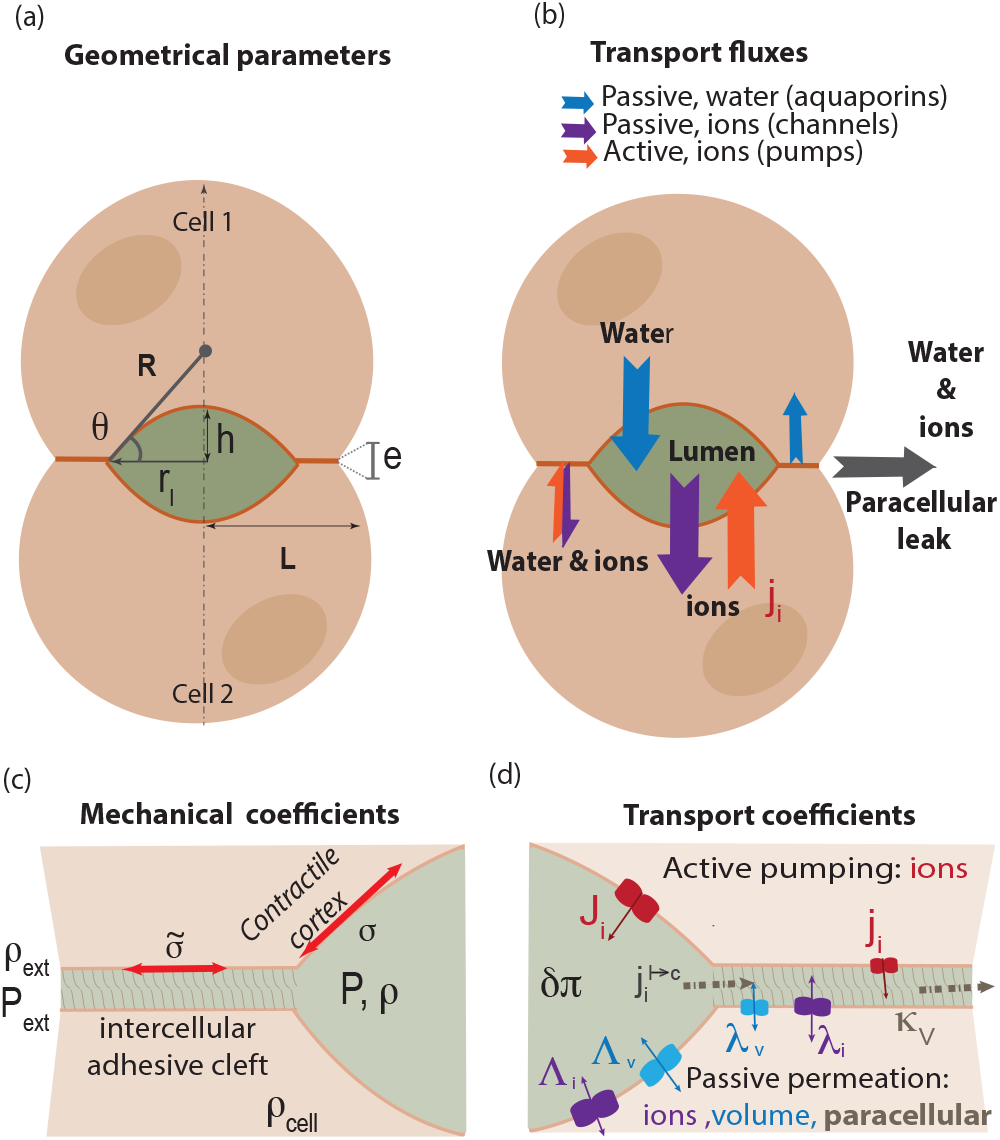
Schematic for lumen at the interface of two adjacent cells.A: Definition of the geometrical parameters of the problem. b: Definition of the active and passive fluxes of Ion and water fluxes across and along the paracellular cleft.C: definition of the mechanical parameters of the problem. Close up on the intercellular cleft region containing adhesive molecules, peptidoglycans and other transmembrane proteins d: Definitions of the transport parameters.

#### Significance Statement

The development of intercellular cavities (lumens) is a ubiquitous mechanism to form complex tissue structures in organisms. The generation of Ciona notochord, the formation of Zebrafish vasculature, or of bile canaliculi between hepatic cells constitute a few examples. Lumen growth is governed by water intake that usually results from the creation of salt concentration difference (osmotic gradients) between the inside and the outside of the lumen. During morphogenesis or in diseases, lumens can also leak due to improper maturation of the cell junctions that seal them. In this paper, we theoretically describe different conditions and dynamical regimes of lumen growth based on the balance of osmotic pressure, fluid intake and paracellular leak.

We study the lumen growth dynamics resulting from the balance between *i* the active and passive ion transport across membranes both in the lumen and in the cleft; *ii* the passive transport of water along transmembrane osmotic and hydrostatic gradients; *iii* the paracellular leakage originating from osmotic gradients and hydrostatic gradients along the cleft; *iv* the mechanical balance controlled by actomyosin contractility. For the sake of simplicity, we considered only one type of anion/cation pair with identical transport properties. These simplified assumptions lead us to consider only ion, water and momentum conservations (i.e, force balance).

### Mechanical balance

In the **lumen** the hydrostatic pressure *δP* is uniform at the time scales considered here. Laplace’s law must be satisfied everywhere across the lumen surface. The force balance in the lumen then reads:

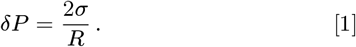

Where *σ* is the cortical tension resulting from the sum of the plasma membrane tension and the active tension of the actin cortex. In general, the effective tension could be inhomogeneous and anisotropic (17). For example, in the late stages of Ciona Notochord lumen growth, or during the tubulation of canaliculi, the departure from a hemispherical shape results in inhomogeneous curvature radii, which is indicative of heterogeneous tension distributions (1, 13, 14). However, here we only consider an homogeneous cortical tension, consistent with the assumption that the lumen shape is a spherical cap. In the **cleft**, Laplace’s law must be modified to account for membrane adhesion (mediated by Cadherin for example (18))

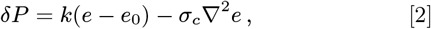

*e*_0_ is the cleft thickness in the absence of a difference in hydrostatic pressure. This is mainly controlled by the cadherin surface density, as well as the repulsive interaction between the membranes. The parameter *k* is an effective elastic modulus that accounts for any deviation of the cleft from *e*_o_, accounting for tension in the cadherins and deformation of the membranes. In SI Appendix (2) we estimate that a few tens of nanometer away from the interfacial region, between the lumen and the cleft, equation 2 results in a homogeneous cleft thickness that hardly deviates from *e*o. In the rest of the paper equation 2 will be replaced by a homogeneous cleft thickness *e*. In the first order approximation *δP* = *k*(*e* **—** *e*_o_).

The force balance at the intersection of the lumen with the cleft is the generalized Young-Dupré equation

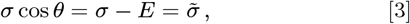

where *θ* is the contact angle (see Fig. 2), *E* is the adhesion energy per unit area, *σ* ˜ corresponds to the “apparent tension” corrected for the adhesion energy. The force balance is thus given by the set of equations 1 and 3.

### Ion conservation

In the **lumen**, ion transport occurs by transmembrane fluxes, as well as by leakage at the lumen edges. The number of ions flowing through the membrane per unit of time and unit of area, has two distinct origins. First, an “active” flux per unit area *J*_*i*_ is generated by pumps and transporters. We assume that the flux has a constant value due to a constant surface density of the relevant pumps.

Ions are also passively transported across trans-membrane channels. In this case, the flux is proportional to the chemical potential difference. It reads 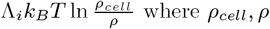 are the ion density in the cell cytoplasm and in the lumen respectively. The transport coefficient Λ_*i*_ is set by the surface density of the relevant channels. By convention all fluxes are positive when ions are secreted into the lumen.

The conservation of the total number of ions, N, in the lumen then reads

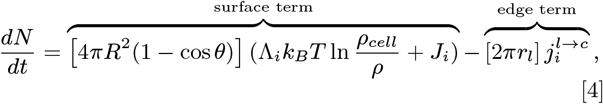

The edge term 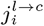 corresponds to the ion flux from the lumen into the cleft. It is determined self consistently by continuity conditions with the expression of the ion flux inside the cleft.

In the **cleft**, the ion density equilibrates within less than a few microseconds across the cleft thickness e (on the order of a few tens of nanometers). Hence, only the ion flux component along the cleft should be considered. The difference in ion concentration in the lumen, as compared to the external medium, generates a diffusive flux 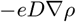 along the ion concentration gradient. D is the diffusion coefficient of ions. We neglect all convective contribution to the flux based on the small dimensions of the cleft. Under these assumptions, and after integration over the constant thickness *e*, the local and time dependent conservation of ions inside the cleft reads

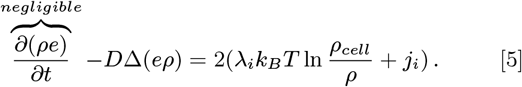

where *λ*_*i*_ is the passive transport coefficient for ions through the membrane into the cleft. *j*_*i*_ is the active pumping of ions. The factor of 2 in the source term accounts for the presence of membranes from both cells. In SI Appendix (2) we show that that the term 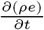 is negligible on the time scale of lumen growth and will further be neglected. 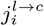 in (Eq. (4) is the solution of equation 5 at *r* = *r*_*l*_.

### Volume conservation

In view of the absence of an active biological transport of water, the change in volume results solely from passive fluxes. Due to water incompressibility, the rate of volume change is proportional to the flux of water. The passive contribution from transmembrane water permeation is proportional to the water chemical potential difference and reads 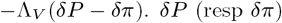 is the difference in hydrostatic (resp osmotic) pressure between the lumen and the cytosol. The surface density of aquaporins determines the transport coefficient Λ_*V*_. The osmotic pressure difference is related to the ion density difference by 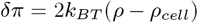. The factor 2 in this expression reflects the equivalent treatment of anions and cations. The conservation of volume in the lumen then reads

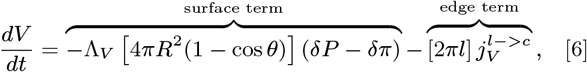

The volume leak 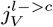 from the lumen into the cleft is determined by continuity of the expression of the volume flux in the cleft at the lumen/cleft interface.

In the cleft, the rapid equilibration of the hydrostatic pressure across the cleft justifies the lubrication approximation to estimate the hydrodynamic contribution of volume change by 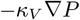. Note that, due to protein crowding at the paracellular cleft, 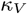 is necessarily smaller than the Poiseuille limit 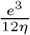 where *η* is the viscosity of the intercellular fluid. The local volume conservation in the cleft then reads

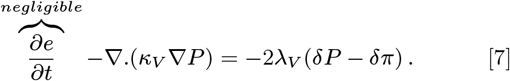

The permeation coefficient *λ*_*V*_ can, in principle, differ in the cleft compared its value in the lumen. For the sake of simplicity, we use the same value. From here on, and for similar reasons as for ion flux, the time derivative of the thickness can be neglected based on the time scale we consider for lumen expansion (see SI Appendix (2)).

### Strategy to solve the equations

The complete set of equations that we solve is provided in SI Appendix (4). To solve the equations, we assume that the parameters of the cytosol and of the external media are constant and homogeneous. We also assume that the variation in ion concentration *δρ,* is small compared to the concentrations themselves. Separating the time scales between lumen dynamics (minutes to hours) and the equilibrium of fluxes in the cleft (sub seconds) simplifies the problem. Cleft equations (3,5,7) are solved in the quasistatic regime. The ion density in the cleft readily stems from Eq. (5). We then use it as a source term in Eq. (7). The solution of Eq. (7) leads to the value of 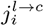, which can in turn can be used in Eq. (4) and Eq. (6). We thus reduce the problem to three coupled equations that we formally solve using Mathematica. SI Appendix Table S1 summarizes the various parameters of the problem and we give their ranges in adimensional and real values in SI Appendix (1).

## Existence of Steady states

At steady state, the dynamical equations above simplify as follows. We name *R*_*s*_, *r*_*s*_ and *θ*_*s*_ the lumen dimensions at steady state.

### Steady state mechanical balance

The Young-Dupré relation takes the simple form

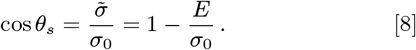

In this expression *σ*_0_ is the steady state tension, and *θ*_*s*_ is thus a constant determined by the tension and adhesion energy at steady state. We take it equal to 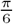 following experimental observations (10).

### Steady state ion conservation

Assuming azimuthal symmetry, the ion conservation in the cleft (eq.5) can be linearized at the first order in polar coordinates as:

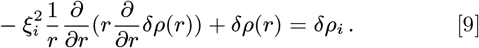

With the continuity equations at the cleft edges being:

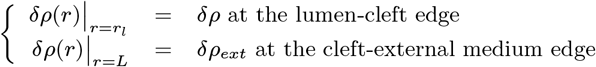

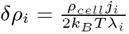 acts as a source term and compares pumping activity to passive ion transport. It corresponds to the ion concentration which would be observed in the cleft if there was a simple balance between pumps and channels. It characterizes the “pumping efficiency”. Note that since *δρ*_*i*_ is a constant, Eq. (9) admits a simple although cumbersome solution in terms of modified Bessel functions, which we give in SI Appendix (3).

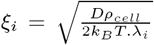 is the typical length over which the ion concentration is screened from the edge effects to reach the constant value set by *δρ*_*i*_. When 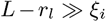 (i.e long cleft and small lumen), the leaks at both edges of the cleft are decoupled from the central part of the cleft the ion density of which only depends on *δρ*_*i*_. Additionally, if *δρ*_*i*_ > *δρ* then, the ion flux 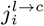 corresponds to an ion source for the lumen. When the lumen is large (i.e *L* — *r*_*l*_ ~ *ξ*_*i*_), the leaks at both edges of the cleft couple to the lumen to create a paracellular concentration gradient. If *δρ > δρ*_*ext*_ the ion flux 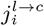 corresponds to a sink for the lumen which takes the simple expression, in the limit 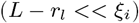:

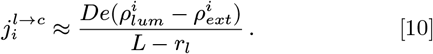

In the **lumen** the ion conservation (4) then simplifies as:

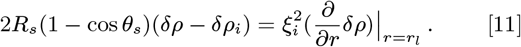

where 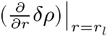 takes the expression derived from the expression *δρ* derived in SI Appendix (3). For the sake of simplicity we assume here that the pump activity in the cleft equals that of the lumen.

### Steady state volume conservation

In the **cleft** Eq. 7 can be simplified in a similar way and writes:

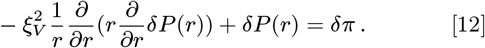

with the continuity of the hydrostatic pressure at both edges imposing:

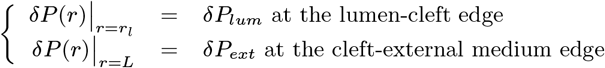

The solution is also tractable analytically (see SI Appendix (3)).

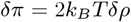 is the source term from osmotic origin. 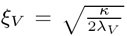 is another screening length, comparing the efficiency of the hydrodynamic leak to aquaporin transport. When 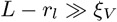, the lumen and the external medium are decoupled. In particular when 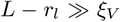 and *ξ*_*i*_ then the hydrostatic pressure in the cleft away from the edges is entirely imposed by the pumps and equals 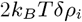.

Whenever the cleft length is longer than both screening lengths, it acts as a volume source for the lumen. In the opposite case (i.e *L — r*_*l*_ ~ *ξ*_*V*_) provided that *P*_*ext*_ < *P*_*lum*_, the cleft contributes to a volume leak out of the lumen that simplifies to

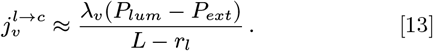

when (*L* — *r*_*l*_ << *ξ*_*V*_).

In the **lumen**, Eq 6 simplifies as

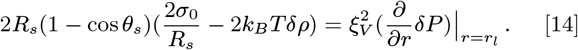

The right hand term is derived from Eq. 12 (see SI Appendix (3)) and taking its value for *r*_*l*_.

This rescaling of the equations reveals that the relevant parameters controlling the lumen are 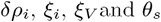. They compare the strength of the various fluxes. They arise from a combination of the more natural parameters 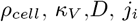 *λ*_*i*_, *λ*_*V*_ and *θ*_*s*_, introduced in the first sections to characterize the fluxes themselves. For all parameter values, the solutions for the steady state lumen radius are qualitatively similar to the one described in Fig.3. For a given leak (characterized by the values of *ξ*_*i*_ and *ξ*_*V*_) there exist a critical value of the ion pumping activity (characterized by *δρ*_*i*_), below which no lumen can exist.

**Fig. 3.**
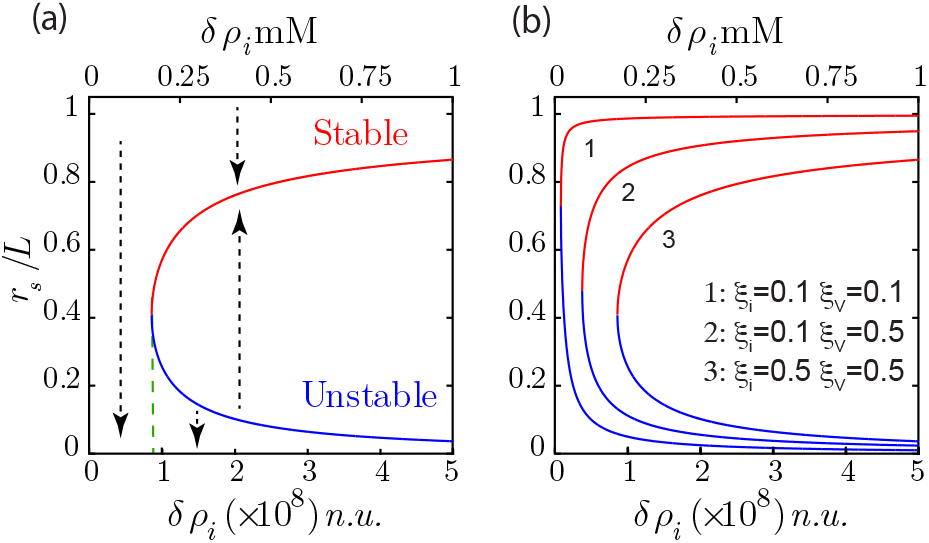
a: The steady state size of the lumen as a function of pumping efficiency displays an unstable and a stable branche represented in blue and red respectively (ξ_*V*_ = *ξ*_*i*_ = 0.5). The dashed arrows represent the direction of variation of lumen radius for any deviation from its steady state value. There is no stable state lumen at low enough pumping efficiency *δρ*_*i*_. Any lumen of any size would shrink off. Above a critical *δρ*_*i*_, any small lumen above the unstable branch will grow to finally reach a larger steady lumen size. b: variation of the steady lumen size as a function of lumen efficiency for different screening lengths *ξ*_*V*_ and *ξ*_*i*_.

Low enough pumping activity cannot compensate the leaks. Independently of its original volume, the lumen shrinks and disappears. When the pump activity is higher, the solution displays two branches. The lower branch is unstable and theoretically corresponds to the creation of a lumen through the nucleation of a small sized cavity inside the cleft. The instability of this solution can be checked directly on dynamical equations, but it can also be understood with the following argument.

Steady state lumens described by lower branches are small (*L* — *r*_*s*_ > ξ_*i*_ and *ξ*_*V*_). A small increase in lumen size leads to a rise in the incoming fluxes, which is due to an increase in lumen surface. However, in this limit the paracellular fluxes are hardly affected by the change in size due to the screening of the leak. Moreover, the osmotic pressure increases, whereas the Laplace term decreases due to tension. Here, the chemical potential balance fails, which leads to further growth. All contributions lead to further volume increase. Although predicted by the model, this solution is likely to be obscured in reality by the more complex biological and molecular organization needed to start lumen formation.

The upper branches correspond to stable solutions for larger lumens (*L* — *r*_*s*_ ~ *ξ*_*i*_ and *ξ*_*V*_). If the lumen grows, the incoming fluxes also grow. However Eq. (10) and Eq. (13) show that in this limit, the paracellular fluxes diverge as the lumen size approaches the size of the junction. This non linear dependence of the paracellular leak in this limit, enables the a stablity of the state. The sensitivity to the edge distance is thus governed by the screening lengths *ξ*_*i*_ and *ξ*_*V*_. Fig.3b shows that small screening lengths (curve 1) result in stable lumens spanning practically the whole cell-cell contact for all pumping activities. Conversely, large screening lengths (curve 3) confine lumens to smaller sizes above a critical pumping activity. One could thus speculate that the ability of lumens from adjacent cell pairs to merge is determined by their ability to reach the cell edges, and is hence controlled by the leak properties of the paracellular cleft.

## Lumen dynamics

The balance between different fluxes not only determines the steady states of the lumen, but also affects lumen dynamics. Fig.1c-d shows that lumen growth can be either monotonous or pulsatile, depending on pumping efficiency. Our model suggests that changing the balance between leaks and ion secretion can induce a transition between both behaviors. The periodicity of the experimental pulsations are of the order of tens of minutes. Consequently, we assume a quasi-static mechanical equilibrium in the cleft. We solve equations 4-2 as described in SI Appendix (5). The time dependent variables of the problem are the radius of curvature *R*(*t*), the contact angle *θ*(*t*) and the difference of ion concentration in the lumen with respect to the cytosol *δρ*(*t*). The lumen shape and volumes can be deduced by simple geometric relations. The cortical tension *σ* must account for the lumen expansion. In situations where the change per unit time of relative cortex area becomes "large", then one must account for a viscous term as a dominant contribution to the periluminal stress. This results in an areal strain rate dependent effective tension. A characteristic time *τ*_*c*_ delineates these two behaviors. In an active gel description of the cortex the effective tension can be written as (19):

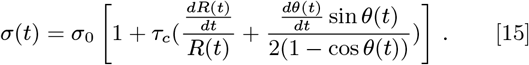

where the quantity 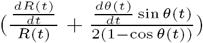 is a measure of the deformation rate, which we take to be equal to the relative time variation of the lumen area. The static value of the tension *σ*_0_ is set by imposing a value of 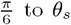 to *θ*_*s*_. All other coefficients are assumed constant. The dynamical equations are expressed in SI Appendix (4).

To exemplify the type of behavior predicted by the model, we fixed the screening length to *ξ*_*V*_ = 0.49, and *ξ*_*i*_ = 0.50 and we solved the dynamical equation at different values of the pumping efficiency *δρ*_*i*_. We set the initial conditions for the lumen height *R(t), θ(t), δρ(t), σ(t)* just above the unstable branch of the lumen steady state (SI Appendix Table 2). In our model this would correspond to a lumen growing from its nucleation size. However the final behavior of the dynamics does not depend on initial conditions. Fig. 4 shows that at lower pumping efficiency, the steady state of the lumen in reached monotonically with a mild overshoot in the contact angle and lumen height. At larger pumping efficiency, the steady state is reached after damped oscillations. At large pumping efficiency the oscillations are sustained. An animation of lumen dynamics in each scenario can be found in Supplementary Videos 1-3. The existence of the oscillations originates from the nonlinearity of the equations, in particular from the divergence of the leak close to the contact edge. However we could not trace one specific parameter alone that was primarily responsible for setting the behavior. In SI Appendix (3) we derive an analytical solution in the transition regime in the limit for large enough lumens 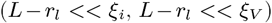 and for small deviations from steady state values of the variables. In the simplified equations, terms analogous to inertia, friction and force could be introduced (respectively a,b and c in SI Appendix (3)); their expressions intricately involve all model parameters. However, the cross over limits between the different dynamic behaviors is set by the parameter *τ*_*c*_, which reflects the dependence of cortical tension on strain rate. Using a constant tension, our numerical solutions do not show any oscillatory behavior within the physiological range of the parameters we explored.

**Fig. 4.**
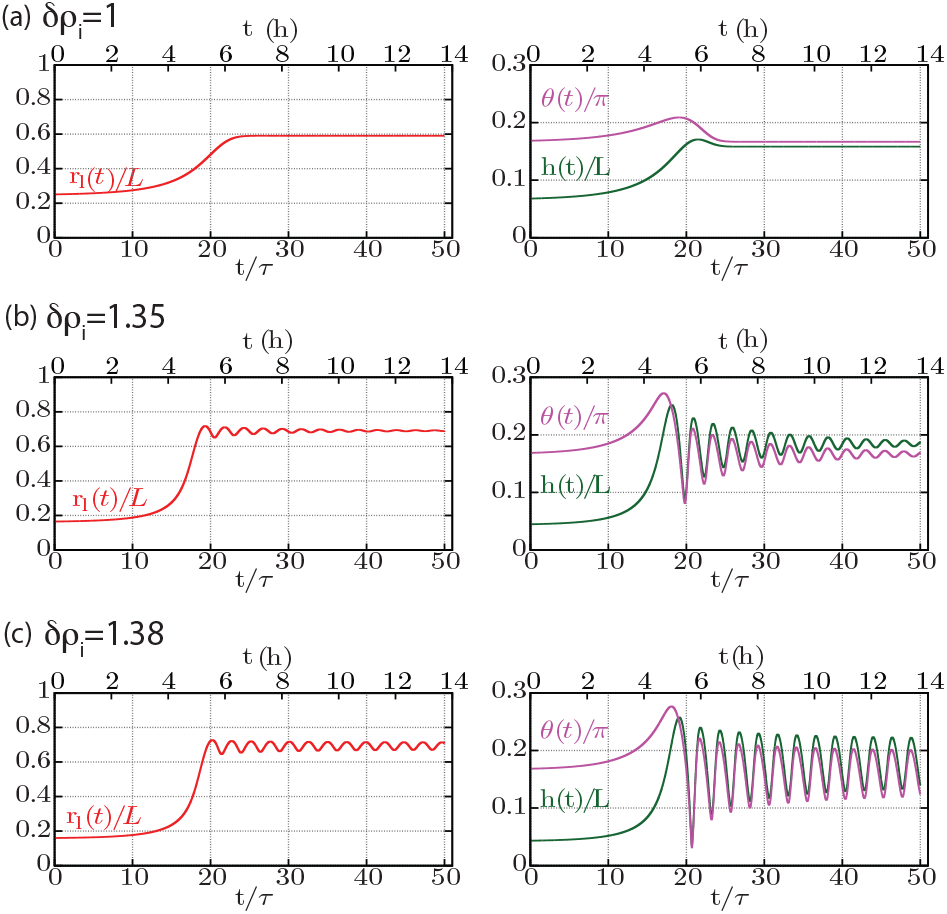
Dynamical behavior of the normalized lumen height *h*(*t*)/*L*, junctional extension *r*_*l*_*(t)/L* and angle *θ(t)/π* are shown as a function of normalized time *t/τ* (lower abscissa) and time in hours (upper abscissa), where *τ* = 2 **x** 10^−8^ *n.u*. is the cortex time (assumed 1000 s). Changing pump efficiency *δρi* shows three different characteristic behavior- (a) Overdamped evolution towards steady state at *δρi* = 1.0 x 10^8^ *n.u*. (b) Underdamped evolution towards steady state at *δρi* = 1.35 x 10^8^ *n.u*. and (c) Sustained oscillations *δρi* = 1.38 x 10^8^*n.u.*. The numerics has been obtained for values of 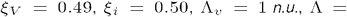 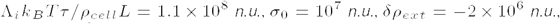, and *ρ*_*cell*_ = 10^9^ *n.u.*.

We then calculated the time variation of the lumen concentration (Fig. 5). In all cases the concentration of the lumen decreases as the lumen grows. It oscillates in phase opposition with the lumen radius in the oscillatory regime. Note however that the total amount of ions *δρ* × *V* increases with the lumen size. The cortical tension varies during the formation of the lumen, increases during the growth phase, and equals *σ*_o_ for the steady states. It oscillates in phase with the lumen radius in the oscillatory case. The inner hydrostatic pressure of the lumen calculated from Laplace’s law decreases as the lumen grows and oscillates in phase opposition with the lumen radius in the oscillatory regime. Our model thus predicts that as the lumen grows the effective periluminal tension grows due to an induced viscous stress. It is qualitatively different from a mechanosensitive feed back that would lead to an active reinforcement of the cortex. Additionally, as the lumen grows the inner pressure decreases. This is the opposite of the "Starling’s law" like interpretation of a lumen growing under an increasing inner pressure, leading to a final contraction that expels the inner fluid. Whereas this later scenario is possible in fully sealed lumen, our model demonstrates that the same dynamical behavior can also be recapitulated in leaking lumens.

**Fig. 5.**
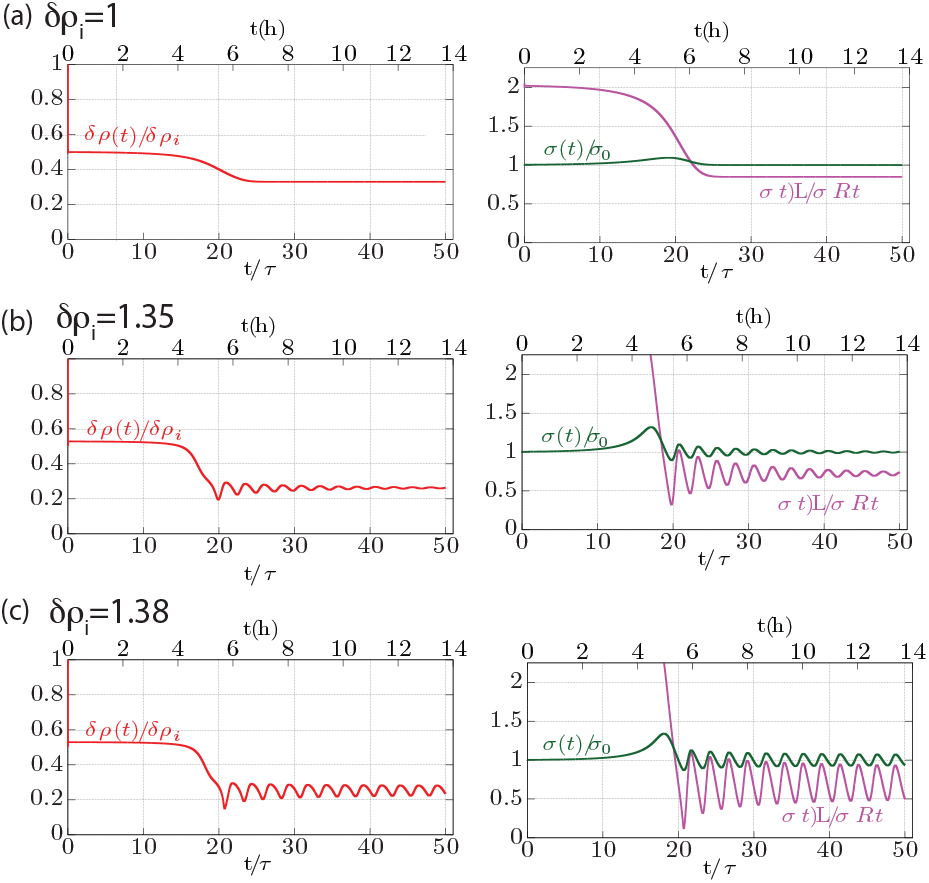
Dynamical behavior of the normalized lumen ion-density *δρ*(*t*)*/δρ*_*i*_, lumen tension *σ*(*t*)/*σ*_0_, and hydrostatic pressure 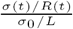 for different pump efficiency *δρ*_*i*_ are shown as a function of time *t*/*τ* (lower abscissa) and time in hours (upper abscissa), where *τ* = 2 × 10^−8^ *n.u*. is the cortex time (assumed 1000 s). Changing pump activity shows three different characteristic behavior- (a) Monotonous over-damped evolution towards steady state at *δρ*_*i*_ = 1.0 × 10^8^ *n.u*. (b) Underdamped evolution towards steady state at *δρ*_*i*_ = 1.35 × 10^8^ *n.u*. and (c) Sustained oscillations *δρ*_*i*_ = 1.38 x 10^8^ *n.u.*. All parameters used for obtaining the numerics are the same as those mentioned in Fig. 4.

## Discussion

The situation of a cavity with constant ion secretion and a fixed cortical tension is intrinsically unstable. A steady state can only be achieved upon three non exclusive conditions: size or time dependent cortical tension, size or time dependent ion secretion, and/or leaks. The two first conditions are likely to involve specific biological feedback. The incidence of leaks is far less intuitive to understand. The model we propose quantitatively explores the effect of paracellular leakage in the case of intercellular lumen formation. We account for the specific dependence of the leak on the dimensions of the paracellular cleft, and we show that, in the case of a bicellular lumens, the leak can play a critical role in controlling lumen size, dynamics and composition. The model provides a good qualitative agreement with the experimental phenotypes of canaliculi. An important prediction of the model is the existence of screening lengths *ξ*_*i*_, *ξ*_*v*_. The screening lengths compare longitudinal fluxes along the cleft that are mediated by osmotic potential differences and hydrostatic pressure, to the transmembrane fluxes that occur orthogonal to the cleft and are mediated by channels. When transmembrane transport outweighs paracellular transport, the screening lengths are small. Curve 1 on Figure 3 shows that in this case the lumen can grow close to the edges (*r*_*s*_ ~ *L*). In contrast, in the case of a large screening length (curve 3) the lumen hardly reaches the cell edge independently of pump activity. The lumen composition, i.e its ion concentration, is also affected by the screening length values. Figure 6a shows that when the distance of the lumen to the cell edge is larger than the screening length the luminal ion concentration is of the same order as *δρ*_*i*_; the equilibrium value for a close lumen. As the lumen grows towards the contact edges, paracellular leaks increase, leading to a decrease in ion density, and hence, of the osmotic pressure as well as hydrostatic pressure. However, Figure 6b, shows that the osmotic pressure decreases considerably less than the hydrostatic pressure. This results in lumens with a much higher ion concentration than what is needed to balance Laplace pressure, should the lumen be closed. Our simplifying assumptions minimize the specific biological details that have yet to be accounted for to perform a quantitative comparison with experimental data. In particular, tight junctions act as diffusive barriers for different classes of ions across claudin pores(20, 21). For the sake of simplicity we account for their activity as a steady factor included in the hydrodynamic resistance of the paracellular cleft. As the tight junctions mature their contribution to the paracellular leak might become dominant over the simple evaluation, which is based on a hydrodynamic process. In particular, ion flux selectivity, which enhanced junction stability and mechanosensitivity of tight junctions, may then play a role in the homeostasis of lumens.

**Fig. 6.**
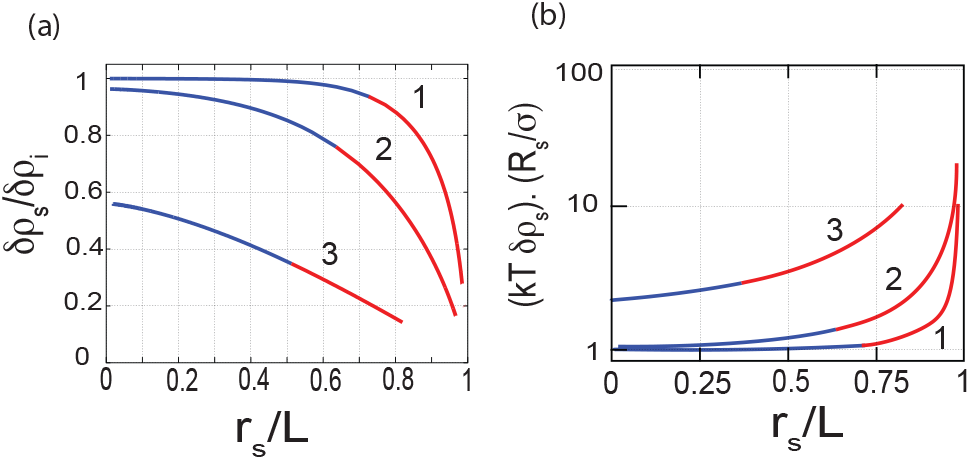
a: Comparison of steady state ion density in the lumens of various sizes with the expected concentration (*δρ*_*i*_). *ξ*_*i*_ = 0.1 for all curves. For curves (1) ξ_*v*_ = 0.1, Curves (2) *ξ*_*v*_ = 0.2, Curves (3) *ξ*_*v*_ = 0.5. b: comparison of the lumen osmotic pressure to the Laplace pressure as a function of lumen size for different screening lengths.

We also show that a time dependent cortical tension is necessary to create an oscillatory behavior. In our model, the origin of cortical tension reinforcement stems from cortex dynamics. As previously mentioned, mechanosensitive mechanisms might reinforce cortex contractility by increasing the actomyosin activity in a stress dependent manner. However, as shown in Figure. 5 the hydrostatic pressure decreases as the lumen grows, and it is not clear where the mechanosensing reinforcement of the cortex would come from within the frame of this model. Although lipid trafficking by endo and exocytosis (1) is important for lumen growth, our model indirectly accounts for it as a non limiting factor of the lumen expansion. Assuming a non limiting rate supply of lipids by vesicular transport, their contribution to cortical tension and thus lumen morphology is negligible. We also do not account for vesicular export of bile in cholestasis cases corresponding to a liver specific problem that would reduce the generality of our description. We indeed propose that the leak dependent growth of lumens can be extended to understand, at the tissue scale, the direction of growth of the cavities. In the case described here, the lumen edge can only asymptotically reach the contact edge due to the divergence of the paracellular leak when *r*_*l*_ approaches *L*. Consider now a single lumen with equal pumping efficiency but embedded in a group of cells rather than a cell doublet. One can qualitatively assume that the resistance to paracellular flux will depend on the total length of paracellular cleft between the lumen edge and the external medium. *L* would then be much larger than the actual size of a single cell-cell contact. In such a case, our model would predict that the lumen radius can extend further than a single cell length and consequently could bridge with other adjacent lumens. Maintaining the same assumptions, the problem of lumen now depends on the structure of the tissue. This more intricate study lies beyond the scope of this work understood as a foundation more elaborate analyses in the future.

